# Evaluating Methods of Updating Training Data in Long-Term Genomewide Selection

**DOI:** 10.1101/087163

**Authors:** Jeffrey L. Neyhart, Tyler Tiede, Aaron J. Lorenz, Kevin P. Smith

## Abstract

Genomewide selection is hailed for its ability to facilitate greater genetic gains per unit time. Over breeding cycles, the requisite linkage disequilibrium (LD) between quantitative trait loci (QTL) and markers is expected to change as a result of recombination, selection, and drift, leading to a decay in prediction accuracy. Previous research has identified the need to update the training population using data that may capture new LD generated over breeding cycles, however optimal methods of updating have not been explored. In a barley (*Hordeum vulgare* L.) breeding simulation experiment, we examined prediction accuracy and response to selection when updating the training population each cycle with the best predicted lines, the worst predicted lines, both the best and worst predicted lines, random lines, criterion-selected lines, or no lines. In the short-term, we found that updating with the best predicted lines or the best and worst predicted lines resulted in high prediction accuracy and genetic gain, but in the long-term, all methods (besides not updating) performed similarly. We also examined the impact of including all data in the training population or only the most recent data. Though patterns among update methods were similar, using a smaller, but more recent training population provided a slight advantage in prediction accuracy and genetic gain. In an actual breeding program, a breeder might desire to gather phenotypic data on lines predicted to be the best, perhaps to evaluate possible cultivars. Therefore, our results suggest that an optimal method of updating the training population is also very practical.

## INTRODUCTION

The improvement of populations in plant breeding through recurrent selection may benefit tremendously from genomewide selection. Of particular worth are the high accuracies and shortened breeding cycles of genomewide selection, which allow for greater genetic gains per unit time (Bernardo and Yu 2007; Heffner *et al*. 2009; Lorenz *et al*. 2011). While genomewide selection has already been employed in established breeding programs for major cultivated species (e.g. Asoro *et al*. 2013; Beyene *et al*. 2015; Sallam *et al*. 2015), this tool also has broad appeal across other species. For instance, breeding programs for tree or perennial crops with long generation times could find utility in making selections before the plants are mature enough to phenotype. Additionally, orphan, undomesticated, or unimproved crops may benefit from rapid breeding progress. Indeed, researchers have already investigated the use of genomewide selection in species such as apple (*Malus* x *domestica*; Kumar *et al*. 2012), *Eucalyptus* (Resende *et al*. 2012), oil palm (*Elaeis guineensis* Jacq.; Cros *et al*. 2015), and intermediate wheatgrass (*Thinopyrum intermedium* (Host) Barkworth & D.R. Dewey; Zhang *et al*. 2016). The population improvement necessary in newly established breeding programs, regardless of species, may be expedited through genomewide selection.

Of course, the aforementioned advantages of genomewide selection depend on maintaining sufficient genetic gain. This requires accurate predictions of the genotypic value of selection candidates based on markers located throughout the genome (Meuwissen *et al*. 2001). Accurate predictions depend on reliable phenotypic measurements and sufficient marker data on a training population. Genomewide marker coverage that captures genomic relationships between individuals and ensures linkage disequilibrium (LD) between markers and quantitative trait loci (QTL) will lead to higher prediction accuracy, especially when predictions are applied to selection candidates more distantly related to the training population (Habier *et al*. 2007; Lorenz *et al*. 2011). The predicted genotypic values under these conditions will more closely reflect the true genotypic values, and selection can then act to increase the frequency of favorable QTL alleles in a population and shift the mean of a population in a desirable direction.

Characteristics of long-term recurrent selection create impediments to maintaining effective genomewide selection. Over generations, recombination between markers and QTL will cause LD to decay, while selection and drift will potentially act to generate new LD or tighten the LD between closely-linked loci (Hill and Robertson 1968; Lorenz *et al*. 2011). Shifts in the pattern of QTL-marker LD, if not captured, will result in decreased prediction accuracy. This suggests that training populations must be updated during recurrent selection to maintain prediction accuracy, a notion that is indeed supported by studies using simulations and empirical data. Studies exploring simulations of recurrent selection in a clonally-propagated crop (*Eucalyptus*) and an inbreeding small grain (barley [*Hordeum vulgare* L.]) both revealed that the accuracy of genomewide selection was improved by updating the training population with data from previous breeding cycles (Jannink 2010; Denis and Bouvet 2013). Similarly, using empirical data from an advanced-cycle rye (*Secale cereal* L.) breeding program, Auinger *et al*. (2016) found that aggregating training population data over multiple cycles enhanced prediction accuracy. These investigations all demonstrated the benefit of including previous-cycle data into a training population, however they did not test different methods of selecting that data.

Though updating the training population may be required, there are practical considerations in how a breeder selects individuals to fulfill this need. Consider a breeding program employing genomewide recurrent selection in barley. Each year, the breeder must allocate phenotyping resources between testing potential cultivars and population improvement.

Though genomewide selection offers to reduce the overall phenotyping costs of the latter (e.g. through early-generation selection), promising breeding lines will undoubtedly be included in field trials. Under genomewide selection, it seems a breeder must also contend with the composition of their training population, placing emphasis on methods to build or maintain this population that both maximize prediction accuracy and minimize costs.

Given the resource limitations of practical breeding and the importance of the training population, it is fitting that much research has been devoted to the composition and design of such populations. Using data from a North American barley breeding program, Lorenz *et al*. (2012) reported reduced prediction accuracy when the training population and selection candidates belonged to separate subpopulations. Multiple studies have found that a training population that is more closely related to the selection candidates leads to more accurate predictions (Asoro *et al*. 2011; Lorenz and Smith 2015). Other researchers have suggested more explicit criteria to determine the optimal training population for a set of selection candidates. Rincent *et al*. (2012) described training population design based on minimizing the mean prediction error variance (PEV) or maximizing the expected reliability of predictions (i.e. generalized coefficient of determination [CD]). When applied to empirical datasets, several investigations supported using the expected reliability criterion to optimally construct training populations (Rincent *et al*. 2012; Akdemir *et al*. 2015; Isidro *et al*. 2015; Rutkoski *et al*. 2015; Bustos-korts *et al*. 2016). These studies generally explored the construction of training populations from a single set of calibration individuals, therefore, the usefulness of this criterion over multiple breeding cycles to maintain prediction accuracy is unknown.

The objective of this study was to investigate various methods of updating a training population and their impact on genomewide recurrent selection. Using simulations, we envisioned a breeding program implementing genomewide recurrent selection for an inbreeding, small grain species (i.e. barley). Six different training population update methods were compared, along with two scenarios of training population composition. We tracked important variables in breeding, including prediction accuracy, response to selection, and genetic variance. Additionally, we attempted to explain some of our observations using other parameters, including persistence of LD phase and genomic relationship.

## METHODS AND MATERIALS

A barley breeding program employing genomewide selection can realistically complete a breeding cycle in a single year (Figure 1). Following this breeding timeline, our experiment simulates a breeding population undergoing 15 cycles of recurrent genomewide selection.

**Figure 1.**
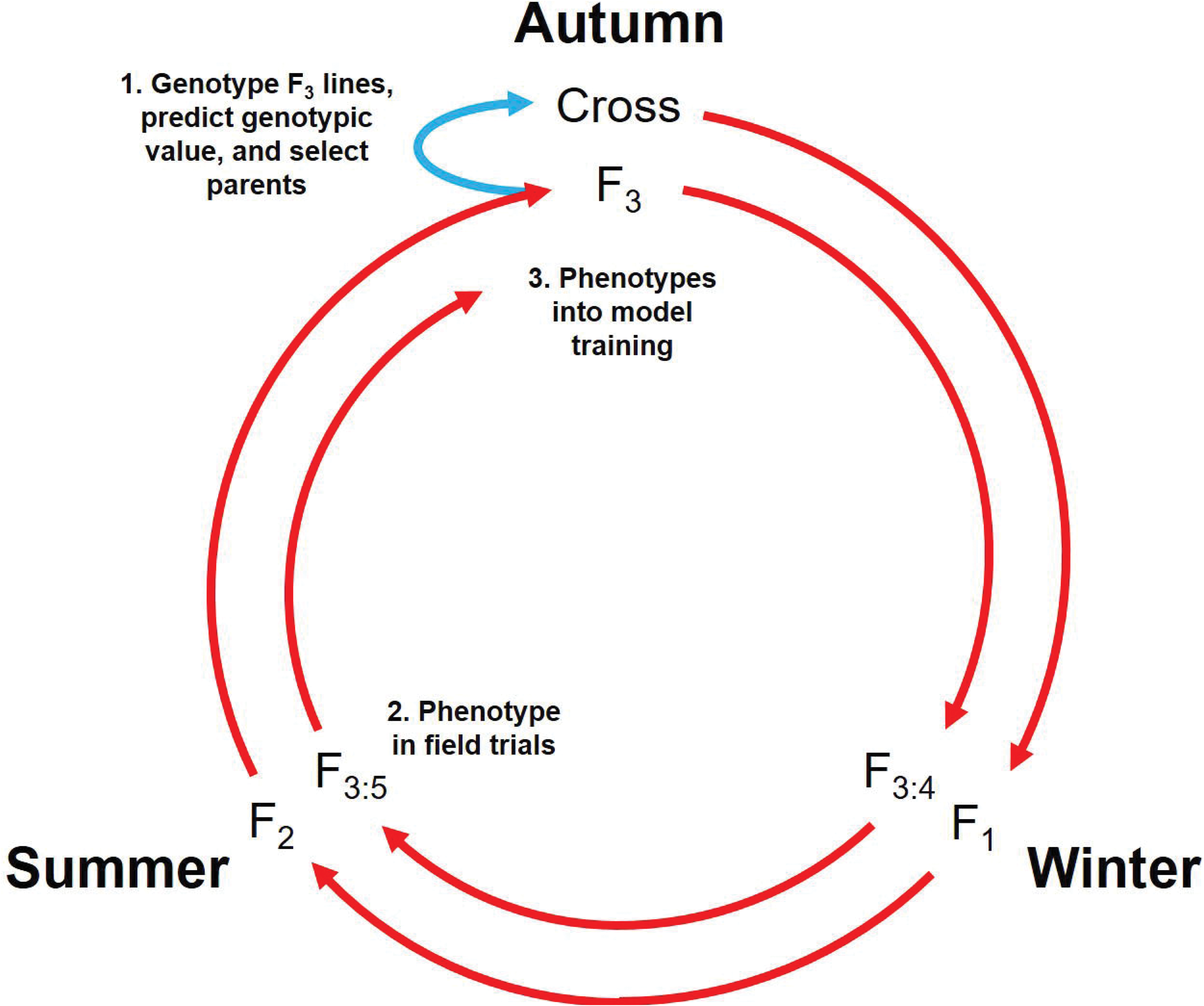
Realistically, a cycle of genomewide recurrent selection in barley may only be one year in length. Crosses are made in the autumn (year*n*) and progeny undergo single-seed descent through the following winter and summer. **1)** At the F_3_ generation during the next autumn (year *n* + 1), lines are genotyped and predicted genotypic value (PGVs) are determined using training data from the previous cycle. These predictions determine the lines to use as parents in the next cycle of crosses (blue arrow). **2)** Predictions are also used to select lines to phenotype in the following summer (year*n* + 2). **3)** This phenotypic information is then incorporated into the training data for the next cycle of predictions and crosses during the subsequent autumn.

To incorporate the observed LD structure in barley breeding populations into our simulations, we used empirical marker data from two North American barley breeding programs: the University of Minnesota (UMN) and North Dakota State University (NDSU). Marker genotypes from 768 six-row spring inbred lines at 3,072 bi-allelic SNP loci were obtained from the Triticeae Toolbox (T3) database (Close *et al*. 2009; Blake *et al*. 2016). The genetic map position of markers was based on the consensus linkage map created by Muñoz-Amatriaín *et al*. (2011). Markers with more than 10% missing data and lines with more than 10% missing data were excluded. Markers were also filtered for redundancy, defined as those located at identical genetic map positions and with identical allele calls. A 0.01 cM interval was forced between markers with non-identical allele calls and shared map positions (i.e. due to low genetic map resolution). We set all heterzyogous genotype calls to missing and imputed missing genotypes using the mode genotype across all samples. This left a set of 764 breeding lines and 1,590 homozygous markers spanning 1,137 cM.

### Genetic Model to Simulate QTL

Each iteration of the simulation was initiated by randomly selecting *L =* 100 SNP loci to become causal QTL, regardless of genetic position or minor allele frequency. Genotypic values for QTL were drawn from a geometric series, as suggested by Lande and Thompson (1990). At the *k*th QTL, the value of the favorable homozygote was *a*^*k*^, the value of the heterozygote was 0, and the value of the unfavorable homozygote was –*a*^*k*^, where *a* = (1 – *L*) / (1 + *L*). The value of the first allele of a QTL was randomly assigned to be favorable or unfavorable. Dominance and epistasis were assumed absent and higher values of the trait were considered favorable. The genotypic value of a given individual was calculated as the sum of the effects of QTL alleles carried by that individual.

Phenotypic values were simulated by adding nongenetic effects to the genotypic values according to the model *y*_*ij*_ = *g*_*i*_ + *e*_*j*_ + *ε*_*ij*_, where *y*_*ij*_ was the phenotypic value of the *i*th individual in the *j*th environment, *g*_*i*_ was the genotypic value of the *i*th individual, *e*_*j*_ was the effect of the *j*th environment, and *ε*_*ij*_ was the residual effect of the *i*th individual in the *j*th environment. Environmental effects were assumed to be samples of a normally-distributed random variable with mean 0 and standard deviation 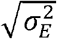, where 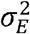 was eight times the variance among genotypic values (i.e. 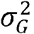) (Bernardo 2014). Residual effects were assumed to be samples of a normally-distributed random variable with mean 0 and standard deviation 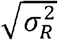, where 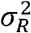 was scaled to achieve a target entry-mean heritability of *h*^*2*^ = 0.5 in the base population. Phenotyping was assumed to take place in three environments with one replication, therefore within-environment variance and genotype-by-environment variance were confounded into 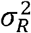. The variance of environmental effects and the variance of residual effects remained unchanged over cycles of selection, allowing the heritability to vary. The mean phenotypic value of each individual over the three environments was used in genomewide prediction.

### Base Population and Cycle 1 of Genomewide Selection

The base population (i.e. cycle 0 training population) consisted of genotypic and simulated phenotypic data on the 764 breeding lines. Based on these simulated phenotypes, the top fifty UMN lines and the top fifty NDSU lines were intermated between breeding programs to generate the cycle 1 population. Specifically, fifty crosses were simulated, using each parent once, and twenty F_3_-derived lines were generated per cross. Gametes were generated following Mendelian laws of segregation, with recombination events simulated according to the genetic map positions of all loci (Muñoz-Amatriaín *et al*. 2011) and assuming no cross-over interference or mutation. Population development resulted in a pool of 1,000 F_3_ selection candidates.

The marker data for the training population and selection candidates consisted of genotypes at all loci except the 100 QTL. This essentially simulated “genotyping” with complete accuracy. Monomorphic markers and those with a minor allele frequency less than 0.03 were removed prior to genomewide prediction. Marker effects were predicted using ridge-regression best linear unbiased prediction (RR-BLUP) according to the model

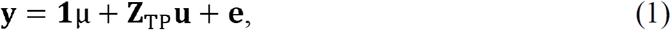
EQN1

where **y** was an *N* × 1 vector of the phenotypic means of *N* training population lines, 1 was a *N* × 1 vector of ones, μ was the grand mean, **Z**_TP_ was a *N* × *m* incidence matrix of training population genotypes for *m* markers, **u** was a *m* × 1 vector of marker effects, and **e** was a *N* × 1 vector of residuals. Elements of **Z**_SC_ were 1 if homozygous for the first allele, -1 if homozygous for the second allele, and 0 if heterozygous. Genotypic values of the F_3_ selection candidates were predicted using the equation 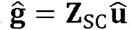 where **ĝ** was a 1,000 × 1 vector of predicted genotypic values, **Z**_SC_ was a 1,000 × *m* incidence matrix of selection candidate genotypes, and 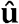 was a *m* × 1 vector of predicted marker effects. Elements of **Z**_SC_ were the same as those in **Z**_TP_.

### Cycles 2 Through 15 of Genomewide Selection

Subsequent cycles of the simulation consisted of three steps: 1) crossing and population development, 2) prediction and selection, and 3) training population updating. These are outlined in the diagram presented in Figure 2. Parents selected in the previous cycle were randomly intermated to form a pool of selection candidates. Again, fifty crosses were simulated and 1,000 F_3_-derived selection candidates were generated. Prior to predictions, we removed monomorphic markers and those with a minor allele frequency less than 0.03 in both the pool of selection candidates and in the training population. Since markers could become monomorphic due to selection or drift, the number of markers used for prediction decreased over breeding cycles. We predicted marker effects by Equation 1, using phenotypic and genotypic data on the training population. These marker effects were then used to predict genotypic values of the 1,000 selection candidates, and those with the top 100 predicted genotypic values were designated as parents for the next cycle. A subset of all selection candidates were then designated as new additions to the training population according to one of the updating methods described below. We simulated phenotypes for these additions and merged the phenotypic and genotypic data to the pool of training population data.

**Figure 2.**
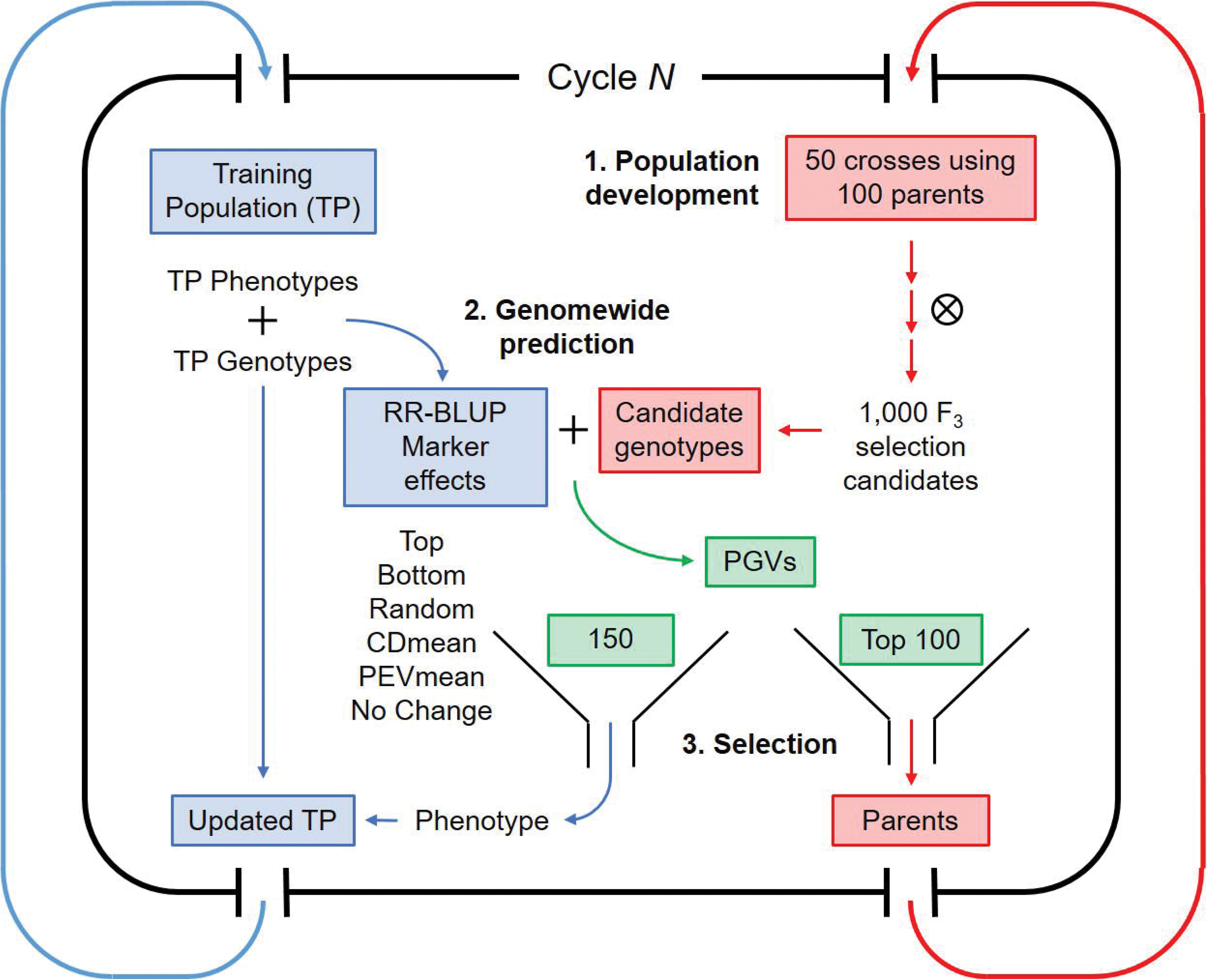
A single breeding cycle in our simulations may be broken down into two main streams. Blue indicates steps involving the training population, and red indicates steps involving crossing and population development. Green indicates the intermediate step of selection. **1)** Fifty crosses are made using 100 randomly intermated parents from the previous cycle. Population development follows and 1,000 selection candidates are genotyped at the F_3_ stage. Concurrently, marker effects are estimated using genotypic and phenotypic data from the training population (TP). **2)** The predicted genotypic values of the selection candidates (PGVs) are used in decision-making. **3)** The 100 selection candidates with the highest predicted genotypic values are selected as parents for the next cycle. Additionally, 150 selection candidates are selected based on the six different update methods. These candidates are “phenotyped”, and phenotypic and genotypic data are added to the pool of training data.

### Methods of updating the training population

Seven different methods of updating the training population were explored in the simulations. For each method, 150 selection candidates from each cycle were selected and added to the training population. These methods are termed “Top,” “Bottom,” “Random,” “PEVmean,” “CDmean,” “Tails,” and “No Change” and are described below. For “Top,” “Bottom,” and “Tails,” selection candidates were ranked based on predicted genotypic value. The 150 selection candidates with the highest (“Top”) or lowest (“Bottom”) values were added to the training population. For the “Tails” method, the 75 selection candidates with the highest values and the 75 selection candidates with the lowest values were added to the training population. For “Random,” a random sample of selection candidates were added to the training population, and for “No Change,” the training population was not updated over breeding cycles.

Two methods involved optimization algorithms previously described by other researchers, specifically “PEVmean” and “CDmean” (Rincent *et al*. 2012). Using only the genotypic data on all individuals, these algorithms aim to create a training population by optimally sampling individuals to be phenotyped in order to predict the value of individuals that would be unphenotyped. Our intention is similar, except that the individuals we sampled to be phenotyped are one cycle removed from the individuals that would be unphenotyped. For PEVmean, selection candidates were chosen to minimize the mean prediction error variance (PEV) of the genotypic values. As described in Rincent *et al*. (2012), the general PEV can be computed using a matrix of contrasts, **C**, between the “unphenotyped” individuals and the mean of the whole population (“phenotyped” and “unphenotyped” individuals). In solving Henderson's (1984) equations, the PEV of any contrast can be computed as

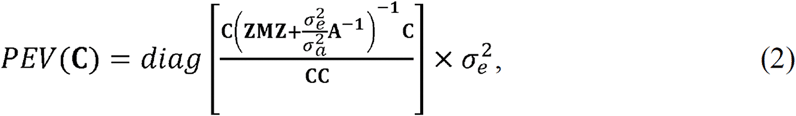
EQN2

where **Z** is an incidence matrix, **M** is an orthogonal projector (Rincent *et al*. 2012), and **A** is the genomic relationship matrix (described below). For the variance of the residuals 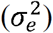, we used the restricted maximum likelihood estimate of 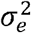 from the RR-BLUP linear model in Equation 1. The additive genetic variance 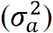 was calculated by multiplying the number of markers, *N*_*m*_, by the restricted maximum likelihood estimate of the variance of marker effects (Bernardo 2014). The PEVmean was then calculated as *PEVmean* = *mean*[*diag*(*PEV*(C))].

Similarly, for “CDmean,” candidates were chosen to maximize the reliability of the predictions, measured as the mean generalized coefficient of determination (CD). This can also be expressed as the expected reliability of the contrasts (Laloe 1993; Rincent *et al*. 2012), computed as

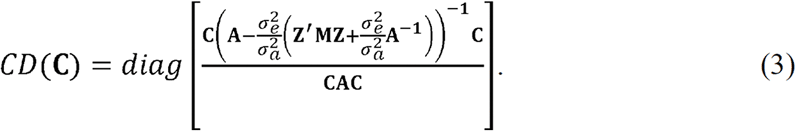
EQN3

The values of 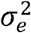 and 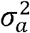 were the same as described for Equation 2. The CDmean was then calculated as *CDmean* = *mean*[*diag*(*CD*(C))]

We implemented an exchange algorithm similar to that described by Rincent *et al*. (2012), with one modification in the designation of individuals to predict and individuals to sample for phenotyping. The situation outlined by Rincent *et al*. (2012) assumes that the genotypic data for the individuals to sample and for the individuals to predict is available concurrently. In our simulation, this is not the case, since phenotyping of the selections in one cycle (cycle*n*) will occur before genotypic data on selection candidates of the next cycle (cycle *n +* 1) becomes available (Figure 1). We therefore chose the 100 parents of the cycle *n +* 1 selection candidates to be a proxy for the unphenotyped individuals, while the entire 1,000 selection candidates (including the parents) constituted the population of individuals to be sampled by the algorithm. To maintain a reasonable computation time, the exchange algorithms were iterated 500 times. Preliminary data showed that a reasonable optimum for either criterion was reached after 500 iterations (data not shown). The PEVmean or CDmean algorithms were used to select individuals from the selection candidates to be included in the training population for the next cycle.

We also considered two scenarios of using the updated training population data. The first scenario represented a situation where a breeder may want to use all available information, and in this case, the training population grew by 150 lines in each cycle. This was termed the “Cumulative” scenario, and over cycles the size of the training population ranged from 764 to 2,864 individuals. In the next scenario, we attempted to control for the effect of training population size by using a “sliding window” of 764 lines along breeding cycles. Specifically, in each cycle the 150 new training population additions from the latest breeding cycle took the place of the 150 training population additions from the earliest breeding cycle. Since the 764 base population lines all constituted cycle 0, these lines were discarded randomly until no base population lines remained in the training population. Afterwards, lines from earlier cycles were discarded as lines from later cycles were added. This was termed the “Window” scenario.

### Variables tracked over breeding cycles

To better interpret the observations in the simulations, we tracked a number of additional variables, including persistence of LD phase, mean realized additive genomic relationship, prediction accuracy, genetic variance, mean genotypic value, inbreeding coefficient, and the frequency of QTL and marker alleles.

The genetic variance in each cycle was calculated as the variance among the genotypic values of the selection candidates. Prediction accuracy was measured by computing the correlation between the predicted genotypic values of the selection candidates and their true genotypic values.

We measured the LD between QTL and markers as such: for each and every polymorphic QTL in a given population (i.e. the training population or the selection candidates), we computed the correlation between that QTL and each and every polymorphic marker in the genome. We calculated persistence of LD phase by first measuring QTL-marker LD in the training population and in the selection candidates. QTL or markers that were not polymorphic in either of these populations were excluded. We then computed the correlation between the measures of QTL-marker LD in the training population and in the selection candidates. This metric, also known as the “correlation of *r*,” evaluates whether patterns of QTL-marker LD are similar between two populations. High correlations of *r* indicate that QTL-marker LD phases are consistent, and presumably the predicted marker effects in one population would accurately represent the marker effects in the second population (de Roos *et al*. 2008; Toosi *et al*. 2010).

Additive relationships between lines in the simulation were measured with respect to the base population. Before initiating the simulations, a matrix **P** was calculated as 2(*p*_*i*_ - 0.5) where *p*_*i*_ is the frequency of the second allele at locus *i* in the base population. Additionally, a normalization constant **c** was calculated as 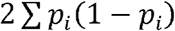. Both calculations are described in VanRaden (2008). To compute additive relationships at any one cycle in the simulation, the genotype matrices (including QTL) of the training population and selection candidates were combined into a matrix **M**. The matrix **P** was subtracted from **M** to obtain matrix **W**. We then calculated the relationship matrix as 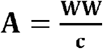. This ensured that the relationship matrix was scaled to reflect the allele frequencies in the base population (VanRaden 2008). We calculated the mean additive relationship as the mean value of the training population-selection candidate combinations. Inbreeding coefficients for each individual were also calculated from this matrix as the diagonal elements minus one.

All simulations were performed in R (version 3.3.1, R Core Team 2016) using the packages *hypred* (version 0.5, Technow 2014) and *rrBLUP* (version 4.4, Endelman, 2011). Each simulation experiment was repeated 250 times. The methods of updating the training population (i.e. “Top,” “Bottom,” “Random,” “CDmean,” “PEVmean,” “Tails,” and “No Change”) each constituted an independent experiment. With the two updating scenarios (i.e. “Window” and “Cumulative”), there were 14 different simulations.

### Data Availability

Simulation scripts, starting marker genotypes, and summarized data are provided in the R package *GSSimTPUpdate*, available from the GitHub repository https://github.com/UMN-BarleyOatSilphium/GSSimTPUpdate. Included is a vignette on how to obtain the marker data from the T3 database.

## RESULTS

### Long-term prediction accuracy

Prediction accuracy (Figure 3, Supplementary Table 1) consistently decreased over cycles of selection for all methods of updating the training population and in both updating scenarios. Within and between scenarios, we observed differences among the update methods in the decay rate of prediction accuracy. A prominent observation was the precipitous decline in accuracy when not updating the training population (i.e. “No Change”). Early in breeding cycles, prediction accuracy for this method was similar to the remaining methods, but by cycle five had decayed beyond the remaining methods. As expected, identical trends were observed for “No Change” in both updating scenarios.

**Figure 3.**
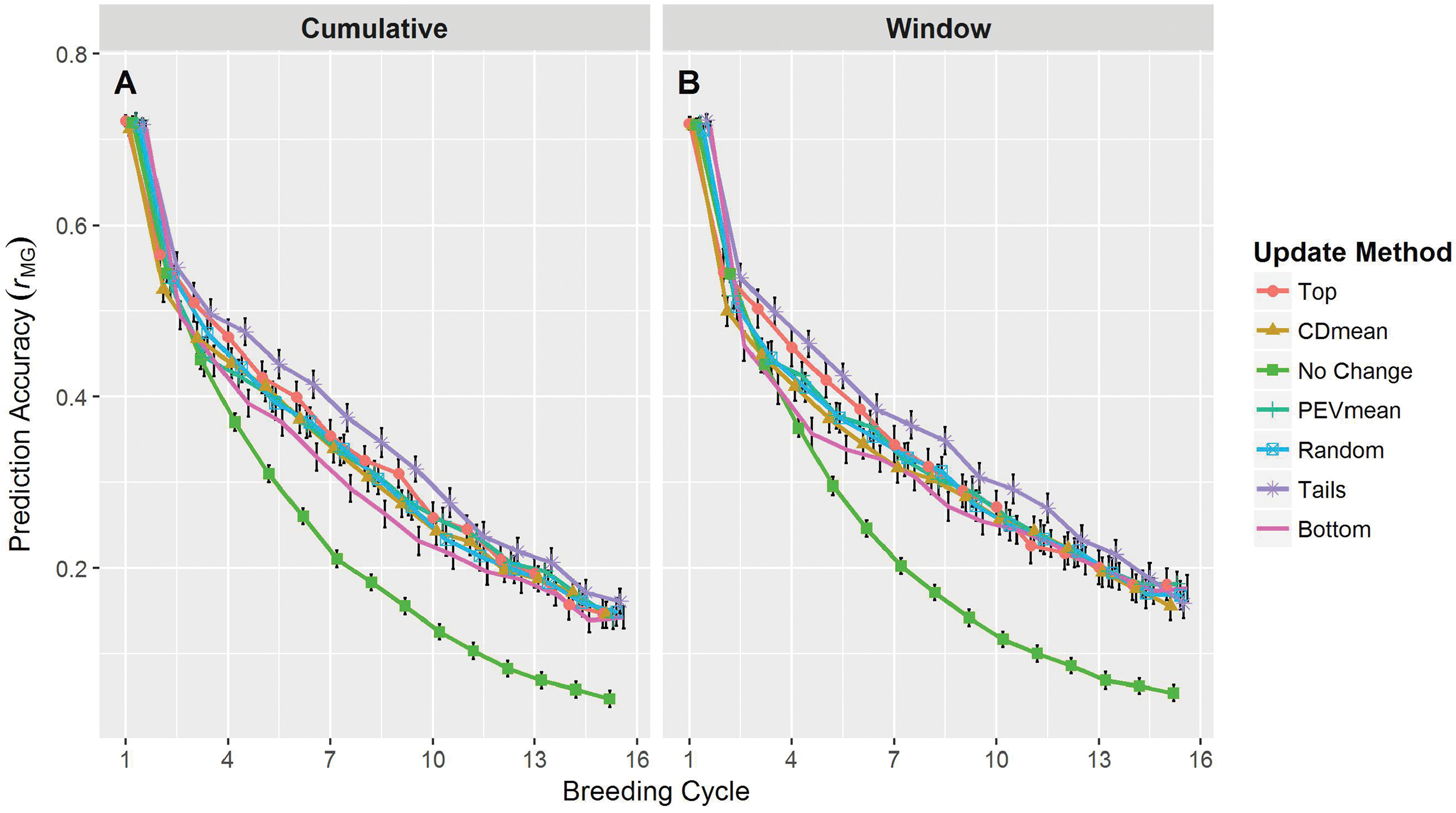
Prediction accuracy over breeding cycles of the simulation. Accuracy was measured as the correlation between the predicted and true genotypic values of the selection candidates. Line colors and point shapes delineate the different methods of updating the training population. Plots are separated into the “Cumulative” (**A**) and “Window” (**B**) updating scenarios. Average values are shown with 95% confidence intervals. To help reduce plot clutter, points for each update method are given a small, consistent jitter along the *x*-axis. Because the plotting jitter may accentuate small differences between updating methods, this data is also provided in **Supplementary Table 1**.

Among methods of actively updating the training population (i.e. excluding “No Change”), differences in prediction accuracy were observed in early cycles, but became increasingly similar in later cycles. The “Top” and “Tails” methods resulted in a non-significant, but noticeable accuracy advantage early on that persisted for several cycles (Figure 3, Supplementary Table 1). On the other hand, the “Bottom” method displayed a noticeable disadvantage that persisted for a similar length of time. The “Random,” “PEVmean,” and “CDmean” methods were highly comparable and yielded accuracies intermediate of the “Top” and “Bottom” methods. By cycle ten, the differences between active methods of updating were negligible. These patterns were observed in both the “Cumulative” and “Window” scenarios.

One noticeable difference between the trends in the “Cumulative” and “Window” scenarios was in the rate of prediction accuracy decay. Among the active methods of updating, the rate of prediction accuracy decay was slightly greater in the “Cumulative” scenario (Figure 3A) compared to the “Window” scenario (Figure 3B). By the fifteenth breeding cycle, the difference in these decay rates amounted to a difference in prediction accuracy of roughly 0.02 – 0.04.

### Genetic variance and response to selection

Genetic variance among the selection candidates (Figure 4A and 4B) similarly decreased across cycles for all training population update methods. For this variable, however, the rank among methods remained more consistent. That is, compared to the remaining update methods, the genetic variance in the “Top” and “Tails” methods was consistently less and the genetic variance in the “Bottom” method was consistently greater. The “Tails” method resulted in slightly higher genetic variance compared to the “Top” method, however this difference was never significant (95% confidence interval). Genetic variance across the “CDmean,” “PEVmean,”, and “Random” methods was very similar within and between scenarios. Not updating the training population resulted in genetic variance similar to “CDmean,” “PEVmean,” and “Random” in early breeding cycles. After seven cycles, however, the loss of genetic variance was abated compared to remaining methods. By the end of the breeding timeline, the genetic variance for “No Change” was noticeably and significantly (95% confidence interval) higher than the remaining methods.

**Figure 4.**
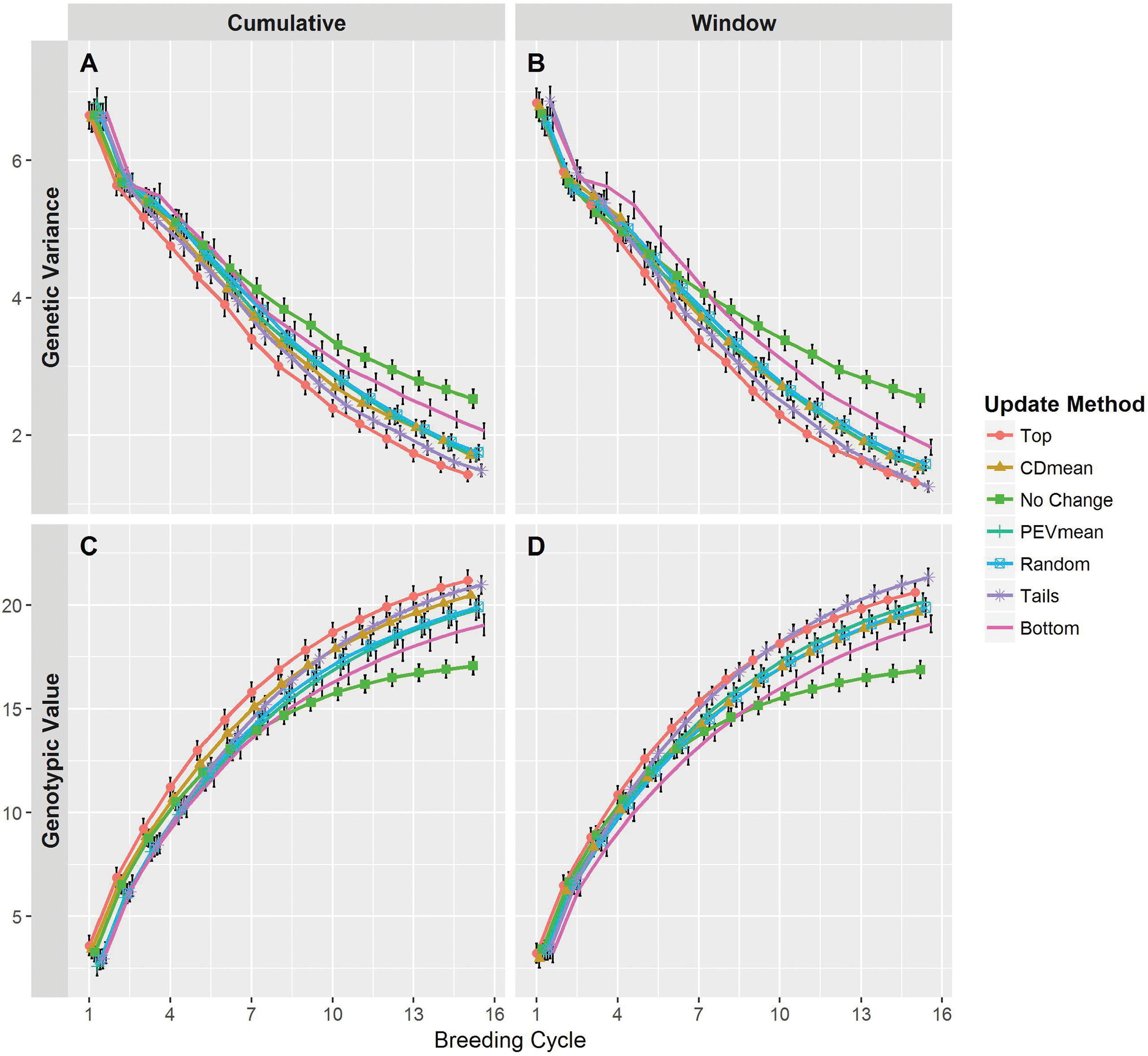
Genetic variance (**A** and **B**) and genotypic values (**C** and **D**) among the selection candidates over breeding cycles of the simulation. Line colors and point shapes delineate the different methods of updating the training population. Plots are separated into the “Cumulative” (**A** and **C**) and “Window” (**B** and **D**) updating scenarios. Average values are shown with 95% confidence intervals. To help reduce plot clutter, points for each update method are given a small, consistent jitter along the *x*-axis.

Overall, the mean genotypic value of the selection candidates (Figure 4C and 4D) displayed a similar, but opposite pattern compared to the genetic variance. Updating the training population by the “Top” or “Tails” methods yielded an advantage in genotypic value, a trend that became more apparent in later breeding cycles. Conversely, the genotypic values under the “Bottom” method ranked lowest among the active updating methods. This disadvantage was often slight and non-significant, especially in the “Cumulative” scenario (Figure 4C). As in the observations of genetic variance, the “CDmean,” “PEVmean,” and “Random” methods responded similarly. Most noticeable was the rapid plateau in genotypic value under the “No Change” method, particularly around the eighth breeding cycle. By the end of the breeding timeline, the “No Change” method appeared to have reached a limit, and although the trajectory of the remaining methods suggested further increases, their trends implied a limit as well (Figure 4C and 4D). Curiously, the “Top” method was generally superior to the “Tails” method in the “Cumulative” scenario, however the opposite was true in the “Window” scenario. In both scenarios, the “Tails” method exhibited a trend suggesting that this method would eventually yield selection candidates with an average genotypic value superior to that of the “Top” method The trends among the remaining training population update methods were similar in both updating scenarios.

### Drivers of prediction accuracy

Average relationship between training population individuals and selection candidate individuals, as measured by marker information, varied among the update methods (Figure 5A and 5B). As expected, the average relationship did not change in either updating scenario when the training population remained unaltered. Across both scenarios, the relationship generally remained highest under the “Top” method, lowest under the “Bottom” method, and intermediate under the “CDmean,” “PEVmean,” “Random,” and “Tails” methods. In the “Cumulative” scenario (Figure 5A), actively updating the training population resulted in a linear increase in average relationship for all methods. Additionally, the different update methods, particularly “Top” and “Bottom,” displayed slight divergence, especially in later breeding cycles. The “Window” scenario (Figure 5B) presented a more sigmoidal trend, eventually resulting in slight convergence in average relationship among active update methods. Interestingly, after cycle 12, the average relationship between the training population and the selection candidates in the “Tails” method remained greater than that in the “Top” method.

**Figure 5.**
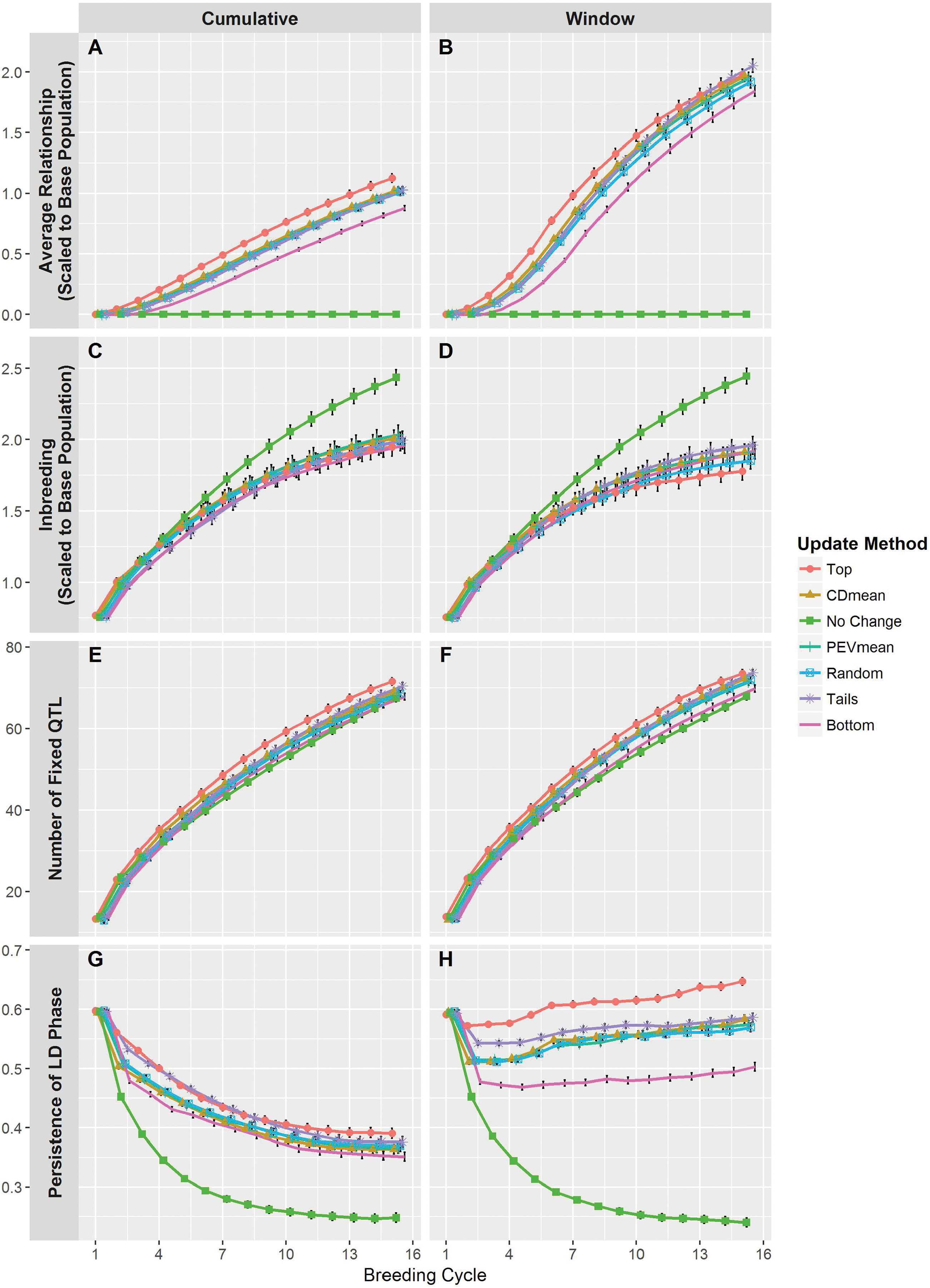
Other variables tracked over the course of the simulations. The average genomic relationship (**A** and **B**) was calculated between the training population and the selection candidates using marker genotypes. Relationships were scaled to reflect the allele frequencies in the base population. The level of inbreeding (**C** and **D**) was measured on the selection candidates and was derived from the relationship matrix described above. The number of QTL fixed for an allele (**E** and **F**) was measured in the selection candidates. Persistence of LD phase (**G** and **H**) was measured as the correlation of *r* between the training population and the selection candidates. Line colors and point shapes delineate the different methods of updating the training population. Plots are separated into the “Cumulative” (**A**, **C**, **E**, and **G**) and “Window” (**B**, **D**, **F**, and **H**) updating scenarios. Average values are shown with 95% confidence intervals. To help reduce plot clutter, points for each update method are given a small, consistent jitter along the *x*- axis.

Generally, we observed a curvilinear increasing trend in the level of inbreeding (Figure 5C and 5D). The “No Change” method performed similarly in the different updating scenarios, but differed markedly from the active updating methods. This method resulted in a more rapid increase in inbreeding, beginning after the fourth breeding cycle. By the end of the breeding timeline, the trend had not yet plateaued and suggested that inbreeding would continue to increase. Considering the active updating methods, there were slight differences in inbreeding trends between the two updating scenarios. In the “Cumulative” scenario (Figure 5C), these methods performed similarly, showing no significant differences. Inbreeding was slightly greater for these methods in this scenario than in the “Window” scenario (Figure 5D). In this case, differences between the updating methods were more apparent. The “Top” method displayed noticeably lower levels of inbreeding, particularly after the eighth breeding cycle. Remaining methods performed similarly between each other.

We noticed consistent trends among methods of updating the training population in the rate of fixation of QTL (Figure 5E and 5F). In both updating scenarios, the “Top” method maintain a higher number of fixed QTL across breeding cycles, followed by the “CDmean,” “PEVmean,” “Tails,” and “Random” methods, which performed similarly, followed by the “Bottom” and “No Change” methods, which also performed similarly. Additionally, we observed that roughly 10% of the QTL became fixed in cycle 1 of the breeding timeline, while by cycle 15 around 70% of the QTL were fixed. There were two slight, noteworthy differences in these trends between the updating scenarios. First, active updating methods generally displayed a higher proportion of fixed QTL in the “Window” scenario (Figure 5E) than in the “Cumulative” scenario (Figure 5F). Second, the degree of separation between the “Top” method and the “CDmean,” “PEVmean,” and “Random” methods appeared greater in the “Cumulative” scenario.

There were marked differences in the persistence of LD phase between the methods of updating the training population within and between the updating scenarios (Figure 5G and 5H). Under the “Cumulative” scenario (Figure 5G), persistence of phase for all update methods declined quickly in initial cycles, but reached equilibrium around the tenth cycle. The “Top” and “Tails” methods maintained the highest degree of persistence across breeding cycles, but the “Tails” method trended closer to the other active update methods by cycle twelve. Furthermore, the initial decay was much lower under the “Top” and “Tails” methods, and the equilibrium point was higher than other methods. Persistence of phase under the “Bottom” method was initially much less than the other active update methods, and although it soon became similar to these methods, it still remained less. The remaining active update methods were quite similar in this scenario.

In comparison, actively updating the training population under the “Window” scenario (Figure 5D) yielded increasing persistence of phase over the course of the breeding timeline. Each of these methods saw a small drop in persistence of phase initially, but after the fifth cycle values began to increase. Interestingly, none of these methods appeared to reach an equilibrium point. The disparity between update methods, especially “Top” and “Bottom,” was highly apparent under this scenario. Conversely, “CDmean,” “PEVmean,” and “Random” resulted in very similar levels of persistence of phase. Finally, the persistence of phase under the “Tails” method was initially intermediate of the “Top” method and the “CDmean,” “PEVmean,” and “Random” methods, however it eventually became more similar to the latter.

Expectedly, the “No Change” method resulted in identical trends in both updating scenarios. In the same way as prediction accuracy, we observed a precipitous, exponential decay in persistence of phase. The trend appeared to reach an equilibrium point at around the same breeding cycle as the active updating methods in the “Cumulative” scenario. However, this equilibrium point was much lower than the others.

## DISCUSSION

### Updating the training population can be simple and effective

We observed similar patterns in prediction accuracy (Figure 3), mean genotypic value (Figure 4C and 4D), and genetic variance (Figure 4A and 4B) among active methods of updating the training population (i.e. excluding “No Change”). The high similarity between these methods suggests that simply including more recent data in the training population provides a marked advantage in improving the breeding population in the long-term. This is encouraging in a practical sense, as any phenotypic information generated on breeding lines, regardless of how they may have been selected, would probably be helpful in preventing severe long-term loss in prediction accuracy.

Although we only tested six active methods of updating the training population, we might expect that any method should outperform doing nothing. Over breeding cycles, including recent genotypic and phenotypic information in the training population helps to capture new LD generated by selection and drift (Hill and Robertson 1968). Older training population lines will of course not provide any information on this new LD, however we may presume most or all selection candidates will share a proportion of this new LD as long as the parents of these lines are not unrelated. Therefore, even the selection candidates most distantly related to those chosen as parents will provide informative training data for the next cycle. In the long-term, we might expect a decrease in the relative importance of how selection candidates are chosen to add to the training population. Over continued cycles of selection in a closed population, parents will become increasingly related (Daetwyler *et al*. 2007), thus the pool of selection candidates will share a greater proportion of the new, informative LD.

Though it appears updating the training population is favorable regardless of method, it is worth pointing out differences in the methods we tested. The “Top” method achieved high prediction accuracy and high mean genotypic value across breeding cycles. These results are not entirely surprising, since the candidates selected to update the training population were mostly those selected as parents for the next cycle (100 of 150). These additions to the training population will be highly related to the selection candidates in the next cycle, and will therefore provide the training population with the most useful information shared through genomic relationships and QTL-marker LD (Lorenz and Smith 2015). Indeed, this is readily apparent in measures of relatedness between the training population and the selection candidates (Figure 5A and 5B) and in measures of persistence of LD phase (Figure 5C and 5D).

With this in mind, it is not surprising that the “Bottom” method delivers the lowest prediction accuracy (Figure 3A and 3B) and lowest mean genotypic value (Figure 4C and 4D), as zero lines added to the training population overlap with the selected parents. This lack of overlap would suggest that QTL-marker LD information in the training additions and that observed in the selection candidates will be in high disagreement. Indeed, we observe that this method produces training populations with the lowest average relationship to the selection candidates (Figure 5A and 5B) and the lowest persistence of LD phase (Figures 5G and 5H).

The “Tails” method, as a combination of the “Top” and “Bottom” method, offers some curious results. Though the prediction accuracy achieved from this method is, for the most part, not significantly different than that of the “Top” method, it is often higher, leading to low genetic variance (Figure 4A and 4B) and high average genotypic value (Figure 4C and 4D). This is in spite of the observation that under the “Tails” method, the average relationship between the training population and selection candidates (Figure 5A and 5B) and persistence of LD phase (Figure 5G and 5H) are roughly equal or lower than in the “Top” method. A possible explanation for this observation could be that this method produces training populations that satisfy different conditions for accurate genomewide predictions. First, 75 of the 150 training population additions overlap with the 100 selected parents. Just as in the “Top” method, these additions will be highly related to the selection candidates of the next cycle and contribute useful QTL-marker LD information. The other 75 additions will presumably be more unrelated to these selection candidates, leading to the intermediate average relationship (Figure 5A and 5B) and often lower persistence of LD phase (Figure 5G and 5H). However, these training population additions may provide information for more reliable predictions. In a study where the training population was a subset of a larger population, Yu *et al*. (2016) found that individuals in the validation population (i.e. selection candidates) with the highest and lowest predicted genotypic values had the greatest upper bound for the reliability of those predictions (Karaman *et al*. 2016). It may be the case in our simulations that the training population additions in the “Tails” method had more reliably-predicted genotypic values. This reliability may have led to better identification of individuals that, when added to the training population, could provide information that more clearly differentiated the effects of QTL alleles, leading to more accurate predictions of marker effects. Thus, the “Tails” method may have taken advantage of both high relatedness and greater genotypic diversity in the training population.

The criterion-based updating methods (“CDmean” and “PEVmean”) performed very similarly to the “Random” method in prediction accuracy (Figures 3A and 3B). This observation is generally in agreement with previous research (Akdemir *et al*. 2015; Isidro *et al*. 2015; Bustos-korts *et al*. 2016) and may be related to the size of the training population used in our simulations. In several examples in these studies, the prediction accuracy of a randomly selected training population was similar to that of a training population selected by the CDmean or PEVmean criteria, particularly at larger sizes of the training population. While these investigations examined training populations ranging from 25 to 300 individuals, our simulations looked at much larger training populations, ranging from 764 to 2,864 individuals. It may be, then, that as the size of the training population becomes sufficiently large, the performance of the CDmean and PEVmean criteria becomes more similar to a random sampling. This, of course, does not suggest that these criteria have no use in selecting training populations. If these criteria are in fact superior in smaller training populations, they may be advantageous when performing genomewide selection on a trait that is expensive or low-throughput to phenotype.

It is worth addressing the continued loss in prediction accuracy in all updating methods and in both updating scenarios. This occurs even as two known components of prediction accuracy, persistence of LD phase and genomic relationship (de Roos *et al*. 2008; Toosi *et al*. 2010; Lorenz *et al*. 2011; Lorenz and Smith 2015; Sallam *et al*. 2015) stabilize or increase. The primary reason for these observations is undoubtedly the reduction in heritability as genetic variance declines over cycles (Figures 4A and 4B). Since residual variance remains constant, the phenotypic data measured on lines becomes increasingly uncorrelated with the true genotypic value (Bernardo and Yu 2007; Bernardo 2010). Thus, the data included in the training population will not capture the effects of QTL alleles, decreasing the accuracy of predicted marker effects. A second potential contributor is the fixation of marker loci over cycles. Since monomorphic markers are removed prior to model training, fewer markers will be used in later cycles. Indeed, by cycle 7, on average 55% of the original markers are used, and by cycle 15 this drops to 30% (data not shown). Though previous studies have stated the benefit of greater marker density (Combs and Bernardo 2013), many others have noted diminishing returns (Lorenzana and Bernardo 2009; Heffner *et al*. 2011; Lorenz *et al*. 2012). Reasonably high marker densities were maintained in our simulations, so this is likely not a strong driver of the decay in prediction accuracy.

The performance of the “Top” method suggests a simple procedure to optimize genomewide selection in an applied breeding program. Our results indicate that a breeder may prevent severe loss of prediction accuracy in recurrent selection by updating the training population to include information on lines that would be selected anyway. Ultimately, this method should be more cost effective than the others. A breeder would likely desire to evaluate selected parents in field trials, perhaps for variety development or to gather phenotypic data to accompany predicted genotypic values. The “Top” method provides an advantage here, as the number of additional lines to phenotype for updating the training population is minimal. The breeder can use this information for dual purposes, using phenotypic data to build a more accurate training dataset while making informed decisions on potential variety selections.

Although the “Tails” method led to slightly greater prediction accuracy than the “Top” method, there are at least three reasons why it may not be the most practical method. First, the difference in prediction accuracy between these methods was generally not significant (Supplementary Table 1). Second, the overlap between training population additions and candidates that would be prioritized for phenotyping by the breeder (i.e. parents and superior lines) is lower, and therefore, third, because of this lack of overlap, the breeder would expend costly resources on phenotyping lines that may not provide any utility outside of model training for genomewide selection.

Encouragingly, empirical data in a barley breeding program supports the “Top” method in enhancing prediction accuracy. Over a few cycles of recurrent genomewide selection for yield and deoxynivalenol content (a mycotoxin produced by the fungal pathogen *Fusarium graminearum* Schwabe.), Tiede (2017, *in prep.*) found that updating the training population improved prediction accuracy. Specifically, including data only on lines selected for favorable predicted genotypic values in previous cycles enhanced the prediction accuracy in subsequent cycles. This method was superior to a random selection of lines and was often superior to a selection based on criteria optimization.

### Not updating the training population is unfavorable

It is quite apparent from our simulations that in the long-term, not updating the training population is highly unfavorable. Prediction accuracy decreases rapidly in this case (Figure 3A and 3B), and as a consequence, response to selection also collapses, leading to the observed plateau in genotypic value (Figure 4C and 4D). Here selection is acting on non-genetic noise, preventing the mean genetic value in the population from changing.

The genetic composition of the breeding populations underscores the negative consequences of leaving the training population unaltered. Although genetic variance appears to be preserved in the long-term (Figure 4A and 4B), considering the decrease in accuracy and the plateau in genotypic value, this may be due to a larger number of QTL that remain segregating. We do indeed observe this (Figure 5E and 5F), but given the similarity in the number of fixed QTL under the “No Change” method and that under the remaining methods, we may also surmise that a greater proportion of QTL are becoming fixed for unfavorable alleles. We also observe alarming levels of inbreeding among the selection candidates when not updating the training population (Figure 5C and 5D). This result is not surprising, since previous theory and simulations into genomewide selection show that more accurate predictions better capture the Mendelian sampling term (i.e. within-family variance), preventing high rates of inbreeding (Daetwyler *et al*. 2007; Jannink 2010). Although higher inbreeding does not reduce genetic variance, it invariably will reduce the number of usable, polymorphic markers. Collectively, this suggests that continued genomewide selection without updating the training population will impose a lower selection limit on population improvement.

The results of our simulations indicate that severe consequences of not updating the training population were delayed until later cycles. Although prediction accuracy declines very rapidly (Figure 3), mean genotypic value and genetic variance track closely with the other updating methods (Figure 4). It is not until the fifth cycle or later that the impact of an unaltered training population is readily apparent. This can be encouraging in practical breeding scenarios. For instance, in a new breeding program, the stock of germplasm with phenotypic data may be low, and it may be several cycles before enough individual are tested to add to the training population. One may also consider a crop where the time between making a cross and gathering phenotypic data on the progeny is long. Several cycles of selection could be performed before data is available to update the training population. Our results suggest that the same training population could be used for a small number of cycles without serious detriment.

### A smaller and more recent training population may provide long-term advantages

We observed non-significant, but noticeable differences in prediction accuracy, mean genotypic value, and genetic variance between the “Cumulative” and “Window” updating scenarios. In the short-term, prediction accuracy was slightly greater under the “Cumulative” scenario for most of the active updating methods, particular the “Top” method (Figure 3A). However, in the long-term, prediction accuracy was higher when the training population consisted of only more recent data (i.e. the “Window” scenario). Although the trends in genotypic value suggest that the “Cumulative” scenario is slightly advantageous in the short-term, the trend under the “Window” scenario suggested that additional gains may be greater (Figure 4D). Indeed, given the slightly higher prediction accuracy observed at the end of the breeding timeline for this scenario, we would expect response to selection to be greater in the long-term (Bernardo 2010).

In addition to the explanations provided earlier in the discussion, other factors may be responsible for these observations. Most notable are the differences between updating scenarios in genomic relationship (Figure 5A and 5B) and persistence of LD phase (Figure 5G and 5H). Retaining older training data results in lower average relationship between the training population and the selection candidates (Figure 5A). This is not unexpected, since selection candidates in earlier cycles will be increasingly unrelated to those in later cycles. Maintaining a training population with more recent data results in higher average relationship and a higher rate of increasing relationship (Figure 5B). This result corroborates previous research demonstrating higher prediction accuracy when retaining individuals in the training population that are more closely related to the selection candidates (Lorenz and Smith 2015).

Perhaps most drastic are the differences in persistence of LD phase between updating scenarios. A training population with older data (i.e. “Cumulative”) results in decayed persistence of LD phase (Figure 5G). Over cycles, recombination breaks down LD and training population additions capture new LD. Older training data does not reflect this new LD, decreasing the persistence of phase. The observed stabilization in Figure 5C could be due to new training data capturing as much LD as what is broken down by recombination. Evidence for this may be seen under the “Window” scenario (Figure 5H), where persistence of LD phase increases when actively updating the training population. A training population of only recent data captures the new LD generated by recombination in the previous cycle, but without the uninformative LD present in older training data. In addition, it may be possible that recent training additions capture more of the informative new LD than what is lost through recombination, leading to the observed increase in persistence of phase.

### Simulation considerations

It is important to address the limitations of our simulations, including assumptions that could be violated in a real-life breeding program. First, random mating may be unrealistic, and we might expect a breeder to impose a more sophisticated procedure for parent selection. For instance, mating pairs may be prioritized for complementation of favorable values of multiple traits. Additionally, an individual may be used as a parent over multiple breeding cycles, especially if observed phenotypic values agreed with the predicted genotypic values. More sophisticated methods of parental selection, such as those based on virtual bi-parental populations (Bernardo 2014; Lian *et al*. 2015; Mohammadi *et al*. 2015), may be used. These non-random mating schemes could affect genetic variance or contribute to different patterns of LD, both of which would impact the accuracy of genomewide prediction. However, incorporating such nuances into our simulation would likely rest on additional assumptions and would be intractable to model. Random mating provides a simple approach, and given the recurrent selection scheme, it is a reasonable assumption. Our simulation also made the assumption that the breeding population was closed. This is obviously inaccurate in a practical program, as the exchange and incorporation of new germplasm is common. Realistically, we might expect prediction accuracy to decrease when adding germplasm from different breeding programs or subpopulations to the pool of selection candidates (Lorenz *et al*. 2012). In recurrent selection, however, the objective is to improve a population rapidly, so a closed population may be desirable (Bernardo 2010).

Other assumptions may not reflect biological reality. First, our simulation forced QTL to be bi-allelic, but, as noted by Jannink (2010) and suggested in Buckler *et al*. (2009), many QTL may have multi-allelic genetic architecture. Second, we assumed the processes of mutation and crossover interference were absent, which is, of course, unrealistic.

## Conclusions

In our simulation experiment of recurrent genomewide selection, we confirmed the need to update the training population over breeding cycles. Clearly, the LD between QTL and markers in the base population is decaying, likely as a result of recombination. When new data is not added to the training population, the change in LD is not captured, and prediction accuracy collapses. Among the tested methods of updating the training population, adding the lines predicted to have the greatest genotypic value (i.e. the “Top” method) is the most attractive. The desirability of this method stems not only from the resulting prediction accuracy and response to selection, but also from its simplicity and practicality. A breeder will undoubtedly desire to confirm the predictions of genotypic value with empirical phenotypic data, especially for the most promising lines or those selected to become parents. Updating the training population becomes simple, then, as this new data can be combined with previous training data. This method also facilitates updating the training model every cycle, likely the best option to capture the changes in LD as a result of recombination, selection, and drift. Nevertheless, our experiment leaves room for additional research, including fine-tuning the updating scenarios to choose the most informative training population from a pool of data. Additionally, optimizing other streams in the breeding program deserves research, including methods of selecting markers and parents. Long-term genomewide selection may benefit greatly from such investigations.

## ACKNOWLEGEMENTS

We thank Celeste Falcon and Ahmad Sallam for their contributions to the discussions from which this research originated. We also thank the Minnesota Supercomputing Institute for the high performance computing resources used to conduct the experiments. This research was partially supported by funding from the Minnesota Department of Agriculture, Rahr Malting Company, the United States Wheat and Barley Scab Initiative, and the National Research Initiative or Agriculture and Food Research Initiative Competitive Grants Program (grant no. 2012-67013-19460) from the United States Department of Agriculture National Institute of Food and Agriculture.

